# Cross Atlas Remapping via Optimal Transport (CAROT): Creating connectomes for any atlas when raw data is not available

**DOI:** 10.1101/2022.07.19.500642

**Authors:** Javid Dadashkarimi, Amin Karbasi, Qinghao Liang, Matthew Rosenblatt, Stephanie Noble, Maya Foster, Raimundo Rodriguez, Brendan Adkinson, Jean Ye, Huili Sun, Chris Camp, Michael Farruggia, Link Tejavibulya, Wei Dai, Rongtao Jiang, Angeliki Pollatou, Dustin Scheinost

## Abstract

Open-source, publicly available neuroimaging datasets—whether from large-scale data collection efforts or pooled from multiple smaller studies—offer unprecedented sample sizes and promote generalization efforts. Releasing data can democratize science, increase the replicability of findings, and lead to discoveries. Due to patient privacy and data storage concerns, researchers typically release preprocessed data with the voxelwise time series parcellated into a map of predefined regions, known as an atlas. However, releasing preprocessed data also limits the choices available to the end-user. This is especially true for connectomics, as connectomes created from different atlases are not directly comparable. Since there exist several atlases with no gold standards, it is unrealistic to have processed, open-source data available from all atlases. Together, these limitations directly inhibit the potential benefits of open-source neuroimaging data. To address these limitations, we introduce Cross Atlas Remapping via Optimal Transport (CAROT) to find a mapping between two atlases. This approach allows data processed from one atlas to be directly transformed into a connectome based on another atlas without the need for raw data access. To validate CAROT, we compare reconstructed connectomes against their original counterparts (i.e., connectomes generated directly from an atlas), demonstrate the utility of transformed connectomes in downstream analyses, and show how a connectome-based predictive model can generalize to publicly available data that was processed with different atlases. Overall, CAROT can reconstruct connectomes from an extensive set of atlases—without ever needing the raw data—allowing already processed connectomes to be easily reused in a wide range of analyses while eliminating redundant processing efforts. We share this tool as both source code and as a stand-alone web application (http://carotproject.com/).

## 1. Introduction

A connectome—a matrix describing the connectivity between any pair of brain regions—is a popular approach used to model the brain as a graph-like structure (Sporns et al., 2004; Bassett and Bullmore, 2006; Bullmore and Sporns, 2009). They are created by parcellating the brain into distinct areas using an atlas (i.e., the nodes of a graph) and estimating the connections between these regions (i.e., the edges of a graph). A wide range of works demonstrates the value of connectomics in studying individual differences in brain function (Elliott et al., 2019; Dubois and Adolphs, 2016), associating brain-behavior associations (Sui et al., 2020; Jiang et al., 2019; Beaty et al., 2018), and understanding brain alterations in neuropsychiatric disorders (Yan et al., 2019). Overall, connectomes have high potential as a biomarker of various phenotypic information.

Nevertheless, the need for an atlas to create a connectome hinders comparisons across studies and replication and generalization efforts. Different atlases divide the brain into different regions of varying size and topology. Thus, connectomes created from different atlases are not directly comparable. In other words, simply comparing the results from two independent studies that use different atlases is challenging. Further, several atlases exist with no gold standards (Arslan et al., 2018), and more are being developed yearly. Currently, no solutions exist to extend previous results and potential biomarkers to a connectome generated from a different atlas, limiting the broader use of potential connectome-based biomarkers.

Transforming an existing connectome into one generated from a different atlas would help these efforts and increase the utility of existing connectomes. For example, large-scale projects—like the Human Connectome Project (HCP) (Van Essen et al., 2013), the Adolescent Brain Cognitive Development (ABCD) study (Casey et al., 2018), and the UK Biobank (Sudlow et al., 2015)—share fully processed connectomes. However, the released connectomes for each project are based on different atlases, preventing these datasets from being combined without reprocessing data from thousands of participants. Smaller labs might not have the resources to store and reprocess these data from scratch (Horien et al., 2021). Finally, due to privacy concerns of being able to identify a participant based on unprocessed data, some datasets are only released as fully processed connectomes (Yan et al., 2019). Critically, in this case, it is not possible to go to the data to create connectomes from another atlas. Thus, algorithms to map and transform connectomes have applications for preserving participant privacy and democratizing data access, as well as improving the generalizability of scientific findings.

To this aim, we propose Cross Atlas Remapping via Optimal Transport (CAROT), which uses optimal transport theory, or the mathematics of converting a probability distribution from one set to another, to find an optimal mapping between two atlases. CAROT is designed for functional connectomes based on functional magnetic imaging (fMRI) data. It allows a connectome constructed from one atlas to be directly transformed into a connectome based on a different atlas without needing to access or preprocess the raw data. We define raw data as data in any form other than fully preprocessed timeseries from an atlas, which is the final form of the data used to create a connectome. Fully preprocessed timeseries from an atlas have several benefits over other intermediate forms derived from a connectomic processing pipeline. As these data consist of only 200 – 500 timeseries, they require less storage than voxel-wise or vertexwise preprocessed data in common space (1 – 3 MB compared to 500 – 1000 MB per individual). These data are also not identifiable if privacy concerns exist.

First, in a training sample with fMRI time series data from two different atlases, we find a mapping by solving the Monge–Kantorovich transportation problem (Kantorovich, 1942). Then, by employing this optimal mapping, time series data based on the first atlas (from individuals independent of the training data) can be reconstructed into connectomes based on the second atlas without ever needing to be preprocessed. To validate CAROT, we compare reconstructed connectomes against their original counterparts (i.e., connectomes generated directly from an atlas), demonstrate the utility of transformed connectomes in downstream analyses, and show how a connectome-based predictive model can be generalized to publicly available data preprocessed with different atlases. Overall, CAROT can reconstruct connectomes from an extensive set of atlases—without ever needing the raw data—enabling comparison across connectome-based results from different atlases and the reuse of already processed connectomes in a wide range of downstream analyses.

This work builds upon two conference papers presented at the 2021 and 2022 International Conference on Medical Image Computing and Computer Assisted Intervention (MICCAI) (Dadashkarimi et al., 2021, 2022). The conference papers present our initial results using optimal transport to map and transform connectomes from different atlases. We expand our previous results by presenting an extensive set of validation studies, increasing the number of atlases tested, and sharing this tool as source code and a stand-alone web application (http://carotproject.com/).

## 2. Theory and calculations

### 2.1. Optimal transport

The optimal transport problem solves how to transport resources from one location α to another *β* while minimizing the cost *C* (Tolstoi, 1930; Hitchcock, 1941; Koopmans, 1949; Gangbo and McCann, 1996). It has been used for contrast equalization (Delon, 2004), image matching (Li et al., 2013), image watermarking (Mathon et al., 2014), text classification (Huang et al., 2016), and music transportation (Flamary et al., 2016). Optimal transport is one of the few methods that provides a well-defined distance metric when the supports of the distributions are different. Other mapping approaches, such as Kullback–Leibler divergence, do not make this guarantee.

The original formulation of the optimal transport problem is known as the Monge problem. Assuming we have some resources *x*_1_,‥, *x_n_* in location *α* and some other resources *y*_1_,‥, *y_m_* in location *β*, we specify weight vectors *a* and *b* over these resources and define matrix *C* as a measure of pairwise distances between points *x_i_* ∈ *α* and comparable points 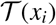. The Monge problem aims to solve the following optimizing problem (Monge, 1781):

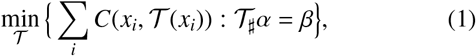

where the push forward operator *#* indicates that mass from *α* moves towards *β* assuming that weights absorbed in 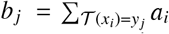. An assignment problem when the number of elements in the measures are not equal is a special case of this problem, where each point in *α* can be assigned to several points in *β*.

As a generalization of the Monge problem, the Kantorvich relaxation solves the mass transportation problem using a probabilistic approach in which the amount of mass located at *x_i_* potentially dispatches to several points in the target (Kantorovich, 1942). An admissible solution for Kantorvich relaxation is defined as 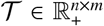 indicating the amount of mass being transferred from location *x_i_* to *y_j_* by 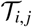:

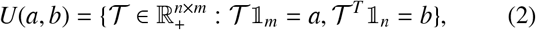

where 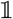 represents a vector of all 1’s. An optimum solution is obtained by solving the following problem for a given 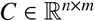 (Rubner et al., 2000):

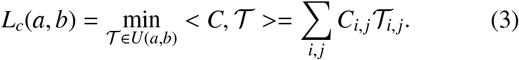

While a unique solution is not guaranteed (Peyré et al., 2019), an optimal solution exists (see proof in Birkhoff (1946); Bertsimas and Tsitsiklis). Kantorovich and Monge problems are equivalent under certain conditions (see proof in Brenier (1991)).

### 2.2. Cross Atlas Remapping via Optimal Transport (CAROT)

CAROT operates by transforming timeseries data from one atlas (labeled the source atlas) into timeseries from an unavailable atlas (labeled the target atlas). This transformation is a spatial mapping between the two atlases. Next, the corresponding functional connectomes can be estimated using standard approaches (e.g., full or partial correlation). Transforming the timeseries data rather than connectomes themselves has two benefits. First, this results in a lower dimensional mapping, which is more robust to estimate. Second, connectomes can be constructed with standard methods (like correlation), guaranteeing properties like semi-positive definite. Directly mapping between connectomes may not guarantee this property.

Formally, let us assume we have training timeseries data consisting of *T* timepoints from the same individuals but from two different atlases (atlas 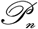 with *n* regions and atlas 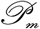 with *m* regions). Additionally, let 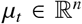 and 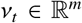 to be the vectorized brain activity at single timepoint *t* based on atlases 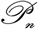 and 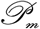, respectively. For a fixed cost matrix 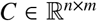, which measures the pairwise distance between regions in 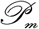 and 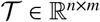, we aim to find a mapping 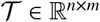 that minimizes transportation cost between *μ_t_* and *ν_t_*:

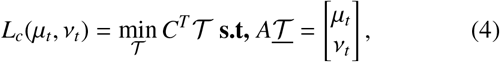

in which 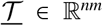 is vectorized version of 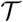 such that the *i* + *n*(*j* − 1)’s element of 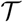 is equal to 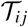 and *A* is defined as:

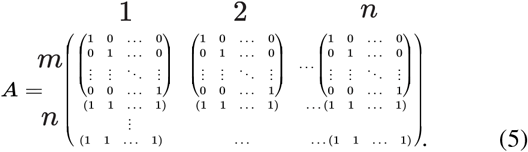

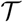 represents the optimal way of transforming the brain activity data from *n* regions into *m* regions. Thus, by applying 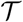 to every timepoint from the timeseries data of the source atlas, we can estimate the timeseries data of the target atlas. As solving this large linear program is computationally hard (Dantzig, 1983), we use the entropy regularization, which gives an approximation solution with complexity of 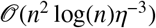 for 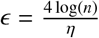 Peyré et al. (2019), and instead solve the following:

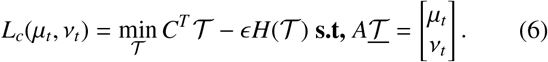

Specifically, we use the Sinkhorn algorithm—an iterative solution for Equation 6 (Altschuler et al., 2017)—to find 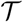. For training data with *S* participants and *K* time points per participant, first, we estimate the optimal mapping 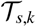, independently, for time point *k* for a given participant *s* using Equation 6. Next, we average 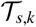 overall time points and participants to produce a single optimal mapping 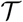 in the training data (*e.g*., 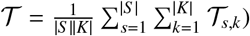.

For the cost matrix *C*, we used a distance metric (labeled functional distance) that is based on the similarity of pairs of timeseries from the different atlases:

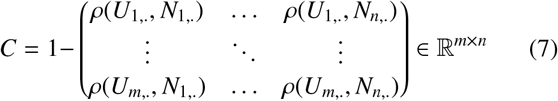

where *U_x_* and *N_x_* are timeseries from 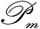 and 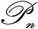 and *ρ*(*U_x_*, *N_y_*) is Spearman correlation between them. To increase a reliable estimation of *C*, we calculate the timeseries correlation independently for each individual in the training data and average over these correlations. Functional distance was used over Euclidean distance between nodes for two mains reasons: (*i*) functional distance does not require having access to the atlas or node locations, which provides greater flexibility should a unknown and unavailable atlas be used, and (*ii*) spatial proximity in the brain does not guarantee similar function. For example, the medial prefrontal nodes of the default mode network are more correlated with nodes in the posterior cingulate cortex than other nodes in the frontal lobe. Nevertheless, we formally compare the performance of functional and Euclidean distances.

## 3. Material and methods

### 3.1. Evaluation datasets

We evaluated CAROT on six prominent functional atlases from the literature using three datasets, the Human Connectome Project (HCP), the REST-Meta-MDD Consortium, and the Yale Low-Resolution Controls Dataset.

#### 3.1.1. Atlases

The Shen atlas (Shen et al., 2013) was created using functional connectivity data from 45 adult participants. The 268-node atlas was constructed using a group-wise spectral clustering algorithm (derived from the N-cut algorithm) and covers the entire cortex, sub-cortex, and cerebellum. The Craddock atlas (Craddock et al., 2012) was created using functional connectivity data from 41 adult participants. The 200-node atlas was constructed using an N-cut algorithm and covers the entire cortex, sub-cortex, and cerebellum. The Schaefer atlas (Schaefer et al., 2018) was created using functional connectivity data from 744 adult participants from the Genomics Superstruct Project (Holmes et al., 2015). The 400-node atlas was constructed using a gradient-weighted Markov Random Field (gwMRF) model, covering only the cortex. The Brainnetome atlas (Fan et al., 2016) was created using structural connectivity data from 40 adult participants from the HCP. The 246-node atlas was constructed using a tractography-based approach and covers the cortex and sub-cortex. The Dosenbach atlas(Dosenbach et al., 2010) was created from meta-analyses of task-related fMRI studies and consists of 160 nodes that cover the cortex, cerebellum, and a few sub-cortical nodes. The Power atlas (Power et al., 2011) was created by combining the meta-analytical approach of the Dosenbach atlas with areal boundary detection based on functional connectivity data. The 264-node atlas covers the cortex, sub-cortex, and cerebellum.

#### 3.1.2. HCP participants

We used behavioral and functional imaging data from this data set as previously described (Gao et al., 2019). We restricted our analyses to those subjects who participated in all nine fMRI conditions (seven tasks, two rest), whose mean frame-to-frame displacement was less than 0.1mm and whose maximum frame-to-frame displacement was less than 0.15mm, and for whom IQ measures were available (n=515; 241 males; ages 22−36+). The HCP minimal preprocessing pipeline was used on these data, which includes artifact removal, motion correction, and registration to common space (Glasser et al., 2013). All subsequent preprocessing was performed in BioImage Suite (Joshi et al., 2011) and included standard preprocessing procedures, including removal of motion-related components of the signal; regression of mean time courses in white matter, cerebrospinal fluid, and gray matter; removal of the linear trend; and low-pass filtering.

#### 3.1.3. REST-meta-MDD

Fully processed data was downloaded from http://rfmri.org/REST-meta-MDD. Full details about the dataset have been previously published elsewhere (Yan et al., 2019). We used data from 21 of the 24 sites. Two sites were removed due to large imbalance between male and female participants (i.e., < 30% male or female; sites 2 and 12). One site was removed as self-reported sex was not provided (site 4). Briefly, the data was processed as follows. First, the initial 10 volumes were discarded, and slice-timing correction was performed. Then, the time series of images for each subject were realigned using a six-parameter linear transformation. After realignment, individual T1-weighted images were co-registered to the mean functional image using a 6 degrees-of-freedom linear transformation without re-sampling and then segmented into gray matter, white matter, and cerebrospinal fluid. Finally, transformations from individual native space to MNI space were computed with the Diffeomorphic Anatomical Registration Through Exponentiated Lie algebra (DARTEL) tool. To minimize head motion confounds, the Friston 24-parameter model was to regressed from the data. Scrubbing (removing time points with FD>0.2mm) was also utilized to verify results using an aggressive head motion control strategy. Other sources of spurious variance (global, white matter, and CSF signals) were also removed from the data through linear regression. Additionally, linear trend were included as a regressor to account for drifts in the blood oxygen level dependent (BOLD) signal. Temporal bandpass filtering (0.01-0.1Hz) was performed on all time series.

#### 3.1.4. Yale participants

In addition, we used resting-state data collected from 100 participants at the Yale School of Medicine. This dataset included 50 females (age=33.3±12.3) and 50 males (age=34.9± 10.1) with eight functional scans (48 minutes total). The dataset and processing details can be found in (Scheinost et al., 2014). Briefly, standard preprocessing procedures were applied to these data. Structural scans were skull stripped using an optimized version of the FMRIB’s Software Library (FSL) pipeline. Slice time and motion correction were performed in SPM8. The remainder of image preprocessing was performed in BioImage Suite. The data was cleaned by regressing nuisance variables (motion parameters, drift terms, and the mean time courses of the white matter, cerebrospinal fluid, and gray matter signals) and band-pass filtering) and was nonlinearly registered to the MNI template.

#### 3.1.5. Generating connectomes

After processing, the Shen, Schaefer, Craddock, Brainnetome, Power, and Dosenbach atlases were applied to the pre-processed fMRI data to create mean timeseries for each node. For each atlas and dataset, connectomes were generated by calculating the Pearson’s correlation between each pair of these mean timeseries and then tasking the fisher transform of these correlations. Connectomes reconstructed by CAROT were also generated using Pearson’s Correlation.

### 3.2. Evaluation overview

We performed several evaluations of CAROT. First, we performed a baseline evaluation of CAROT, investigating the similarity of the original and reconstructed connectomes, the impact of free parameters (e.g., the number of participants used to train CAROT), and the number of available source atlases. Second, we investigate how reconstructed connectomes perform in standard downstream analyses (i.e., do reconstructed connectomes give similar neuroscience results as the original connectomes?). Finally, we present a real-world evaluation of how CAROT can generalize a preexisting connectome-based predictive model when data from the required atlas is unavailable.

### 3.3. Baseline evaluation of CAROT

#### 3.3.1. Similarity between the original and reconstructed connectomes

We compared reconstructed connectomes to their original counterpart using HCP data. We partitioned our data into a 25/75 split, where 25% of the individuals is used to estimate the optimal mapping 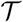 and 75% is used to evaluate the reconstructed connectomes. Reconstructed connectomes were created using single and multiple source atlases. To evaluate the similarity between CAROT reconstructed and original connectomes, the upper triangles of the connectomes were vectorized and correlated with Spearman’s rank correlation.

#### 3.3.2. Evaluation of free parameters

We investigated the sensitivity of CAROT to the number of time points and number of participants used to find the mappings and the value of *ϵ* in the Sinkhorn approximation. Using the same 25/75 split of the HCP participants for training and testing as above, we varied the number of time points used from 100 to 1100 in increments of 100, varied the number of participants from 100 to 515 in increments of 100, and varied ε from 0.01 to 10 in increments of 1.

#### 3.3.3. Extending CAROT for multiple atlases

A vital drawback of the single-source optimal transport is that it relies on a single pair of source and target atlases (i.e., one source atlas and one target atlas), which ignores additional information when multiple source atlases exist. As preprocessed data is often released with timeseries data from multiple atlases (Yan et al., 2016), we investigated using these additional data to better reconstruct connectomes from an unavailable atlases. To incorporate multiple atlases, we applied CAROT to each pair of atlases, transforming timeseries data for each source atlas into the timeseries data for the target atlas. Next, the transformed timeseries data are averaged across all source atlases, and a single connectome for the target atlas is created (Fig. 1). Further, we investigated the impact of using a smaller number of source atlases by only including *k* random source atlases when creating a connectome for the target atlas. This process was repeated with 100 iterations over *k* = 2−6.

**Figure 1:**
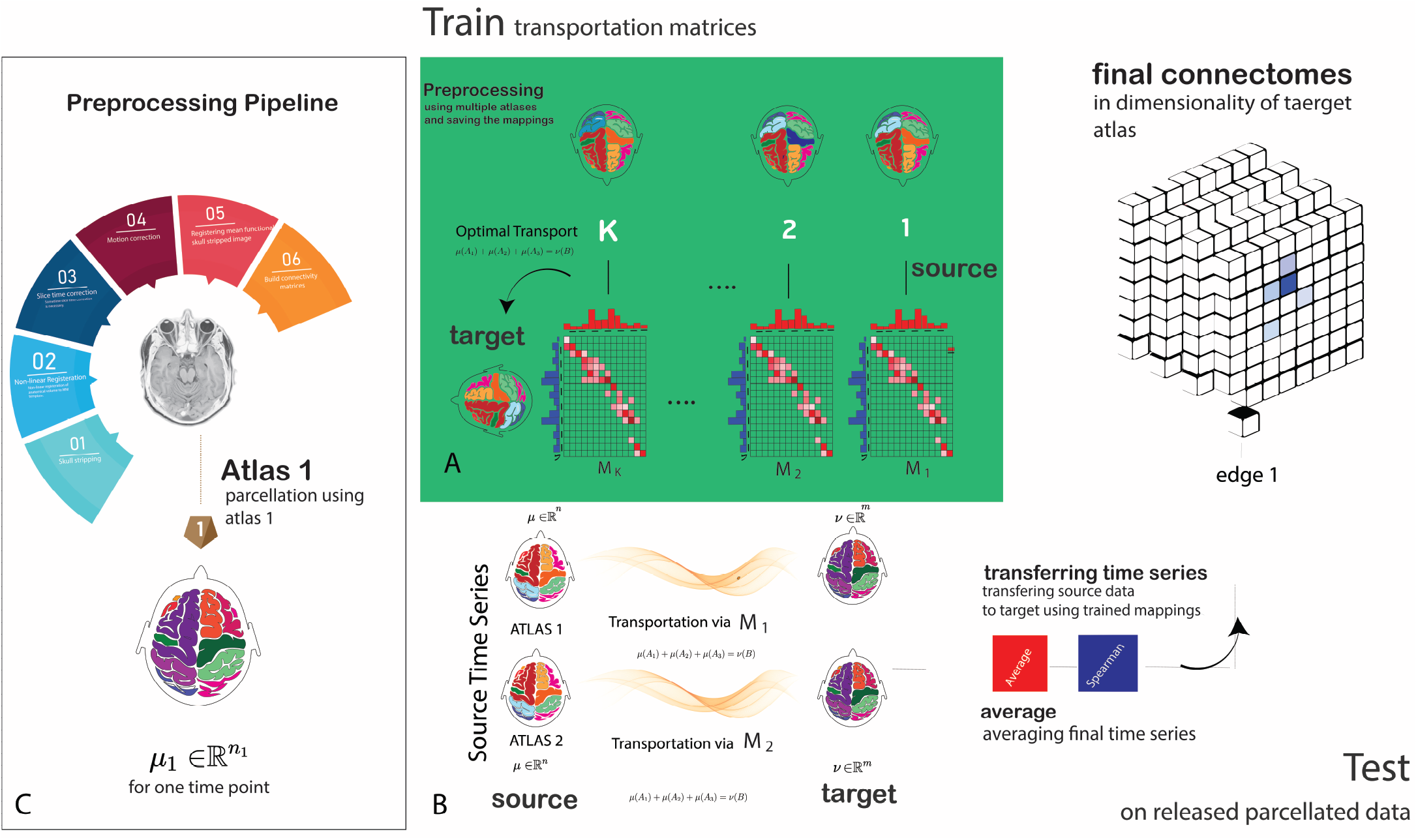
Schematic of CAROT: A) During training, CAROT transforms timeseries fMRI data from multiple source atlases to a target atlas to obtain transportation mappings. Mappings between the source and target atlases are found by employing optimal transport and solving Monge–Kantorovich transportation problem using the Sinkhorn approximation. The solution provides a transformation that maps the brain activity parcellated using the source atlas to brain activity parcellated based on the target atlas. B) During testing, for each pair of source and target atlases and a single time point in the timeseries data, the offline solutions are used, and time series and functional connectomes accordingly will be reconstructed in the desired target atlas. Results from several pairs of source and target atlases can be combined to improve the quality of the final reconstructed connectome. C) A standard image preprocessing pipeline to create functional connectomes.

#### 3.3.4. Generalizing mappings across datasets

We investigated if CAROT mappings trained in one dataset generalize to other datasets. In other words, we tested if CAROT can be trained once (for example, using the HCP) and then be applied to any new datasets without the need to rerun CAROT (for example, the Yale dataset). First, we trained CAROT using only the HCP dataset. Then, we reconstructed connectomes using the Yale dataset using these 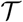’s. Spearman’s rank correlation between the upper triangles of the connectomes was used to assess the similarity between the reconstructed and original connectomes.

### 3.4. Evaluation of downstream analyses

#### 3.4.1. Consistency of aging results

We tested that the reconstructed connectomes produced consistent neuroscience results compared to the original connectomes. First, we used 25% of the HCP participants to train CAROT. Next, using the original and reconstructed connectomes for the remaining 75% of participants, we calculated the association between connectomes and age using mass univariate, edge-wise correlations. Results were threshold at *P* < 0.05, corrected for multiple comparisons using the Network-based Statistic (NBS) (Zalesky et al., 2010). To assess whether the overlap of the significant edges found using the original and reconstructed connectomes was statistically significant (i.e., edge-level), we calculated the probability of the overlap being due to chance using the hypergeometric cumulative distribution:

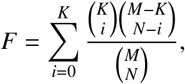

where *F* is the probability of drawing up to *i* of a possible *K* items in *N* drawings without replacement from a group of *M* objects. The p-value for the significance of overlap is then calculated as 1 − *F*. We also assess the similarity of results at the node-level by summing over all significant edges for a node (i.e., the network theory measure—degree) and correlating these maps for results from the original and reconstructed connectomes.

#### 3.4.2. IQ prediction

To show that meaningful brain-phenotype associations are retained in reconstructed connectomes, we used reconstructed connectomes to predict fluid intelligence using connectome-based predictive modeling (CPM) (Shen et al., 2017). We partitioned the HCP dataset into three groupings: *g*_1_, consisting of 25% of the participants; *g*_2_, consisting of 50% of the participants; and, *g*_3_, consisting of the final 25% of the participants. In *g*_1_, we trained CAROT for each source and target atlases pair. We then applied the learned 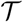 on *g*_2_ and *g*_3_ to estimate connectomes for each target atlas, resulting in nine connectomes for each atlas (seven reconstructed connectomes based on a single source atlas, one reconstructed connectome based on all source atlases, and the original connectome). Finally, we trained a CPM model of fluid intelligence for each set of connectomes using *g*_2_ and tested this model in *g*_3_. Fluid intelligence was quantified using a 24-item version of the Penn Progressive Matrices test. Spearman correlation between observed and predicted values was used to evaluate prediction performance. This procedure was repeated with 100 random splits of the data into three groups.

#### 3.4.3. Identification rate

We investigated if the individual uniqueness of connectomes is retained in reconstructed connectomes by identifying individuals scanned on repeated days (Finn et al., 2015). As mentioned above, we used the HCP data and a 25/75 split to create reconstructed connectomes based on all available source atlases. In an iterative process, one individual’s connectome was selected from the target set and compared against each of the connectivity matrices in the database to find the matrix that was maximally similar. Spearman correlation between the target connectome and each in the database was used to assess similarity. A score of 1 was assigned if the predicted identity matched the true identity, or 0, if it did not. Each target connectome was tested against the database in an independent trial. Connectomes generated from the day 1 resting-state data were used as the target set, and connectomes generated from the day 2 resting-state data were used as the database. We performed this identification procedure for the original and reconstructed connectomes independently. We used permutation testing to generate a null distribution to determine if identification rates were achieved at above-chance levels. Specifically, participants’ identities were randomly shuffled and identification was performed with these shuffled labels. Identification rates obtained using the correct labels were then compared to this null distribution to determine significance.

### 3.5. Real-world evaluation

In this evaluation, we generalize a sex classification model (using 100 adults collected at the Yale School of Medicine and created with the Shen atlas) to the REST-Meta-MDD dataset (Yan et al., 2016), which only provides preprocessed timeseries data from the Dosenbach, Power, and Craddock atlases. First, we trained the sex classification model using the Yale dataset’s resting-state data from 100 individuals (50 males). We trained a *ℓ*_2_-penalized logistic regression model with 10-fold crossvalidation to classify self-reported sex. Then, we used 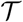 estimated from the HCP to transform the publicly available preprocessed data (i.e., timeseries data from the Dosenbach, Power, and Craddock atlases) from the REST-Meta-MDD dataset into the Shen atlas. Data from each source atlas were combined to create a single connectome based on the Shen atlas for the 1005 (585 females) health controls. Finally, the sex classification model created in the Yale dataset was applied to these reconstructed Shen connectomes.

### 3.6. Data availability

All datasets used in this study are open-source: HCP (ConnectomeDB database, https://db.humanconnectome.org), REST-meta-MDD (http://rfmri.org/REST-meta-MDD), and Yale dataset (http://fcon_1000.projects.nitrc.org/indi/retro/yale_lowres.html). BioImage Suite tools used for processing can be accessed at (https://bioimagesuiteweb.github.io/). CAROT and associated canonical mappings are on GitHub (https://github.com/dadashkarimi/carot). The Python Optimal Transport (POT) toolbox is available at https://pythonot.github.io/.

## 4. Results

### 4.1. Baseline evaluation of CAROT

#### 4.1.1. Reconstructed connectomes are similar to original connectomes

As shown in Table 1, the correlation between the reconstructed connectomes and their original counterparts depends on the atlas pairing, with more similar atlases appearing to have higher correlations. For instance, a strong correlation is seen while transforming data from the Craddock atlas to the Shen atlas (*ρ* = 0.48, *p* < 0.001). Both atlases are based on clustering timeseries fMRI data using variants of the N-cut algorithm. In contrast, a weaker correlation exists for transforming data from the Dosenbach atlas (which was constructed based on a meta-analysis of task activations) to the Shen atlas (*ρ* = 0.24, *p* < 0.001). Finally, similarity between the reconstructed and original connectomes was much lower when using Euclidean distance (Fig. S1).

**Table 1:** Spearman correlation between reconstructed connectomes and original connectomes for each source-target pair. Presented results show as mean ± standard deviation over 100 random splitting the data into training and testing sets.

|  | Shen | Schaefer | Craddock | Brainnetome | Power | Dosenbach |
| --- | --- | --- | --- | --- | --- | --- |
| Shen | | 0.46 $\pm 0.010$ | 0.55 $\pm 0.001$ | 0.42 $\pm 0.003$ | 0.36 $\pm 0.030$ | 0.39 $\pm 0.004$ |
| Schaefer | 0.31 $\pm 0.001$ | | 0.38 $\pm 0.010$ | 0.38 $\pm 0.001$ | 0.33 $\pm 0.010$ | 0.34 $\pm 0.002$ |
| Craddock | 0.48 $\pm 0.010$ | 0.54 $\pm 0.010$ | | 0.51 $\pm 0.003$ | 0.43 $\pm 0.002$ | 0.43 $\pm 0.001$ |
| Brainnetome | 0.17 $\pm 0.003$ | 0.23 $\pm 0.003$ | 0.23 $\pm 0.002$ | | 0.19 $\pm 0.004$ | 0.18 $\pm 0.002$ |
| Power | 0.24 $\pm 0.002$ | 0.35 $\pm 0.003$ | 0.32 $\pm 0.001$ | 0.29 $\pm 0.070$ | | 0.32 $\pm 0.002$ |
| Dosenbach | 0.24 $\pm 0.001$ | 0.32 $\pm 0.003$ | 0.28 $\pm 0.020$ | 0.28 $\pm 0.001$ | 0.28 $\pm 0.010$ | |

#### 4.1.2. Using multiple atlases improves CAROT

Overall, we observed a considerable improvement when including data from multiple source atlas. In every case, using all available data produced more similar connectomes to their original counterparts (all *ρ*′s > 0.50; Fig. 2). For most atlases, explained variance is more than tripled using CAROT with multiple source atlases compared to using a single source atlas. As shown in Fig. 2, while the similarity between reconstructed and original connectomes increases as the number of source atlases increases, strong correlations (e.g., *ρ* > 0.6) can be observed with as little as two or three source atlases, suggesting that a small number of atlases may be sufficient for most applications.

**Figure 2:**
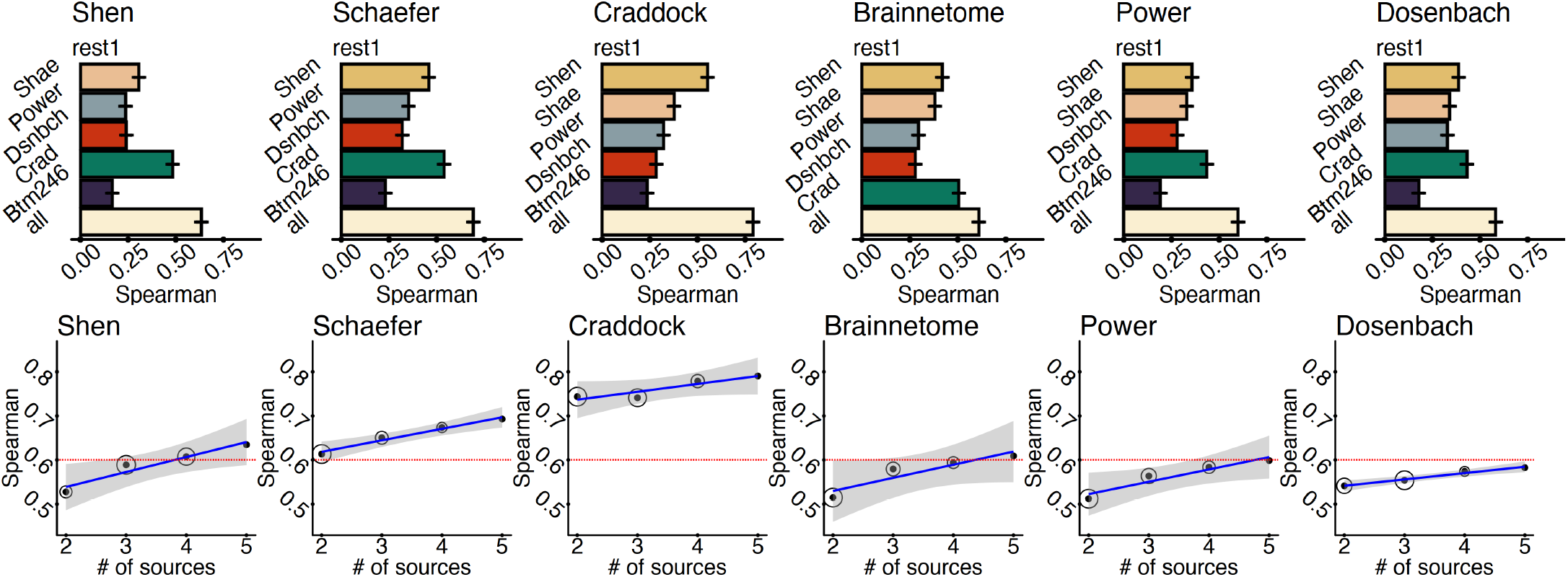
Using multiple source atlases improves the similarity of reconstructed connectomes. A) The Spearman’s rank correlation between the reconstructed connectomes and connectomes generated directly with the target atlases are shown for each pair of source and target atlas as well as reconstructed connectomes using all of the source atlases. Using all source atlases produces higher quality reconstructed connectomes for each target atlases. Error bars are generated from 100 iterations of randomly splitting the data into 25% for training and 75% for testing. B) For each target atlas, increasing the source atlases increases the similarity of reconstructed and original connectomes. For most atlases, a Spearman’s correlation of *ρ* > 0.60 (red line) can be achieved by using fewer than five source atlases (i.e., all available source atlases). Circle size represents the variability of the correlation over 100 iterations of splitting the data into training and testing sets.

#### 4.1.3. CAROT is insensitive to parameter choices

No clear pattern of performance change was observed across the tested parameter range, suggesting that CAROT is not affected the number of frames and participants, and the range of *ϵ*’s (Fig. S2). However, using only 100 participants and 100 time points significant (*p* < 0.05) reduced the processing time from 2,975 s to 467s.

#### 4.1.4. Mappings generalize across datasets

When applying the mapping trained in the HCP dataset to the Yale dataset, we observed a strong correspondence between the reconstructed connectomes and their original counterparts with *ρ’s* > 0.50 (Shen: *ρ* = 0.59; Schaefer: *ρ* = 0.66; Craddock: *ρ* = 0.71; Brainnetome: *ρ* = 0.54; Power: *ρ* = 0.50; Dosenbach: *ρ* = 0.54). Notably, these correlations are in the same range as those observed when applied these mappings to the HCP data (i.e., the same dataset used for training the mappings). Together, these results exhibit that mappings can be trained in one dataset and applied to another.

### 4.2. Evaluation of reconstructed connectomes in downstream analyses

#### 4.2.1. Similar patterns of aging are found with reconstructed connectomes

At the edge-level, the reconstructed connectomes for all atlases produced aging results that significantly overlapped with the results from using the original connectomes (*p* < 0.00001). Similarly, node-level correlations were all significant (*r′s* > 0.60, *p′s* < 0.001). Fig **??** shows a representative example of node-level results between the reconstructed and original connectomes for the Shen atlas.

#### 4.2.2. Reconstructed connectomes predict IQ

In all cases, connectomes reconstructed using all source atlases performed as well in prediction as the original connectomes (Fig 4A). The reconstructed connectomes using all source atlases performed better than the original connectomes for the Schaefer and Power atlases. Similar to other analyses, connectomes reconstructed from a single atlas varied in prediction performance, depending on the combination of source and target atlases.

**Figure 3:**
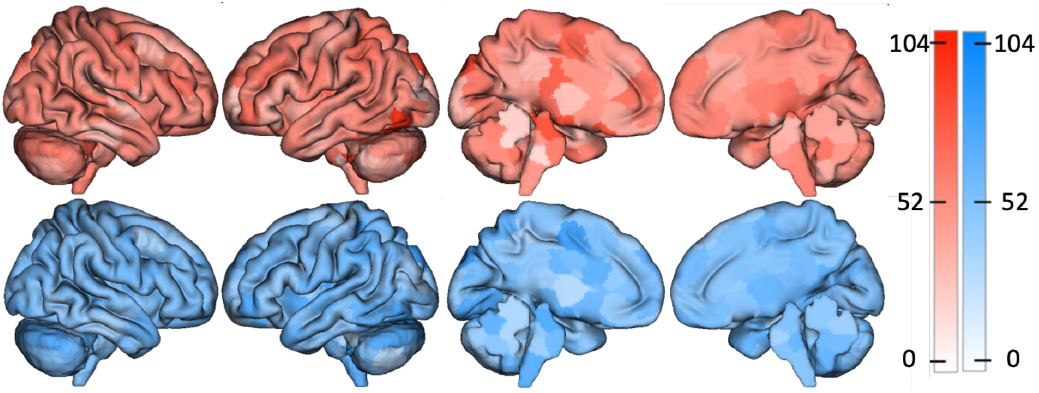
Reconstructed connectomes give similar aging results as the original connectomes. The top row shows the nodes with the largest number of edges significantly associated with age for original connectomes from the HCP created with the Shen atlas. The bottom row shows the same but using reconstructed Shen connectomes. These spatial maps correlate at *r* = 0.61, suggesting that analyses with the reconstructed connectomes produce the same neuroscientific insights as analyses with the original connectomes.

**Figure 4:**
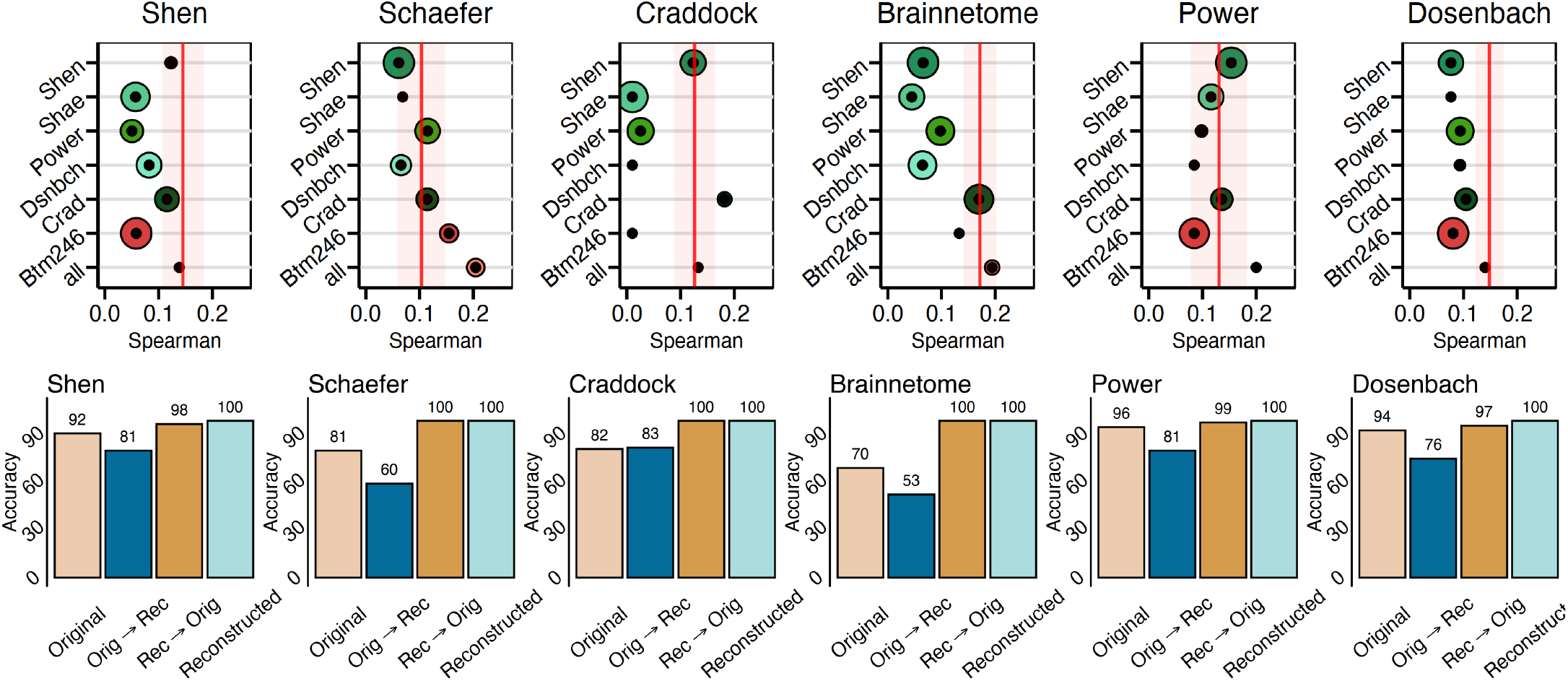
Reconstructed connectomes behave the same as original connectomes in downstream analyses. A) The reconstructed connectomes retain sufficient individual differences to predict IQ using connectome-based predictive modeling. In all cases, reconstructed connectomes based on all available source atlases (bottom circle) predicted IQ as well or better than the original connnectome (red line). Size of circle represents the variance of prediction performance of 100 iteration of 10-fold cross-validation. B) The reconstructed connectomes retain sufficient individual uniqueness to identify individuals using the reconstructed connectomes.

#### 4.2.3. Reconstructed connectomes are unique to an individual

For all analyses, identification of individuals demonstrated a high success rate that were significantly greater than chance (5%; *p* < 0.001; based on permutation testing). (Fig. 4B). Reconstructed connectomes performed slightly bettwer than the originals (original connectomes: mean rate=79%; reconstructed connectomes: mean rate=90%). Overall, these results suggest that the reconstructed connectomes retain similar levels of individual differences as their original connectome counterparts.

### 4.3. CAROT facilitates external validation of connectome-based predictive models

Overall, the sex classification model demonstrated significant classification accuracy in the Yale dataset (Accuracy=60.5% ± 6%; Naive model accuracy=50%; *χ*^2^ = 5.8; *p* = 0.03). Next, the sex classification model preformed significantly better than chance in the REST-Meta-MDD dataset when using the reconstructed connectomes (Accuracy=66.5%; Naive model accuracy=52.3%; *χ*^2^ = 13.9; *p* = 0.0002). To better this result into context, we created connectomes for the Dosenbach, Power, and Craddock atlases in the Yale dataset, created a sex classification model for connectome type, and generalized these models to the REST-Meta-MDD dataset. The generalization accuracy of reconstructed connectomes (Shen: 66.5%) was numerically superior to the generalization accuracies based on original connectomes (Dosenbach: 59.6%, Power: 59.0%, and Craddock: 64.5%), suggesting that using CAROT and reconstructed connectomes perform as well as original connectomes in generalizing a preexisting predictive model.

### 4.4. Software availability and implementation

To facilitate open science and the broader adoption of CAROT, we have created http://carotproject.com/. This web application allows end-users to convert timeseries data from the Shen, Schaefer, Craddock, Brainnetome, Power, and Dosenbach atlases to connectomes for any of the other atlases. As a web application, it works without software installation and across multiple platforms (*e.g*., Windows, Linux, MacOS, Android). The only requirement is a modern web browser, such as Google Chrome. Please note, that any data used on http://carotproject.com/ remains on the local computer and is never uploaded or stored on a remoted server. In addition, we provide the CAROT software and associated canonical mapping as opensource at https://github.com/dadashkarimi/carot/.

Specifically, we provide functionality: *(i)* to generate the cost matrix based on functional distance for timeseries data from two different atlases; *(ii)* to generate the mapping 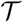 between between two atlases based on the cost matrix defined above; and *(iii*) to convert timeseries data from one or more source atlases to connectomes based on a target atlas. In addition, we provide canonical mappings based on the HCP data to map between every pair of the Shen, Schaefer, Craddock, Brainnetome, Power, and Dosenbach atlases. Based on the results present here, these mappings should work in other datasets, saving researchers the need to regenerate these mappings for themselves. We will look to provide mappings between additional atlases as they become available. CAROT is implemented in Python 3, building on the Python Optimal Transport (POT) toolbox (Flamary and Courty, 2017).

## 5. Discussion and conclusions

Neuroimaging is at a crossroads, facing a need to increase replication efforts and use larger-than-ever samples (Yarkoni, 2009; Szucs and Ioannidis, 2020; Marek et al., 2022). These are tough challenges for functional connectomics, where connectomes created from different atlases are incomparable. As such, processed connectomes or connectomic results from different atlases must be reprocessed from raw data. Here, we introduced and validated CAROT, a method that will allow us to overcome the limitation of not being able to combine connectomes and results from different atlases. CAROT allows functional connectomes from different atlases to be transformed to a common atlas and combined in downstream analyses. CAROT relies on optimal transport to find a frame-to-frame mapping of fMRI time series data used to create functional connectomes for a missing atlas. We show that these reconstructed connectomes are highly similar to the original ones and perform similarly in downstream analyses. Specially, reconstructed connectomes retain sufficient individual differences to predict IQ and uniqueness to identify individuals. Finally, we provide a real-world example of how a connectome-based predictive model (based on the Shen atlas) can be generalized to open source, preprocessed data that was not processed with the Shen atlas.

Critically, the mappings between connectomes are general to the dataset used to created the mappings. As such, a single set of canonical or gold-standard mappings can be trained with one dataset and be distributed to work in new datasets without re-training the mappings. Accordingly, we have released initial mappings based on the HCP data to map between every pair of the Shen, Schaefer, Craddock, Brainnetome, Power, and Dosenbach atlases as part of our software. We hope that CAROT and http://carotproject.com/ will save researchers time and effort by eliminating data reprocessing and increase the ease of performing mega-analysis and external validation efforts.

Across analyses, we show that CAROT produces reconstructed connectomes that achieve similar results in IQ prediction and fingerprinting as connectomes created directly from the data. This observation holds across a range of atlases that differ in their construction and constituent brain regions. While atlas pairs that are more similar in terms of their construction and coverage produced better pair-wise mappings (e.g., the Craddock and Shen atlases were created with N-cut algorithms and cover the cortex, sub-cortex, and cerebellum (Shen et al., 2013; Craddock et al., 2012)), using multiple source atlases is even better. Likely, combining transformed time series averages out the minor idiosyncrasies in the individual mappings between atlas pairs, producing more stable results. Overall, when using multiple source atlases, CAROT is robust to differences between the source and target atlases.

While including all available data generated the most similar connectomes, strong correspondence between reconstructed and original connectomes was observed when as little as 2 or 3 different source atlases were used. Together, this suggests that an exhaustive list of every possible atlas does not need to be released, but that only including a few different atlases could vastly increase the utility of any released preprocessed data. Balancing the utility of released data and the effort to release it is a delicate task. If the data is not in a convenient form for end-users, it will not be used and, if the effort is too high to share data, data will not be shared. We believe that CAROT can help balance these, by increasing the utility of the shared data with only a slight increase in effort for sharing the data.

Given that using multiple source atlases produces more robust results, it encourages future studies to release preprocessed data from a few atlases. Not only does this increase the chances that the needed atlas is available for an end-user, but also better facilitates the use of the data when the needed atlas is unavailable. Some open source datasets already release data from multiple atlases (e.g., REST-Meta-MDD). That CAROT preforms better with multiple altases may be relevant for large-scale projects, like ABCD and UK Biobank, that share raw data and curated releases. Given that these datasets range in the several thousands of participants, curated data from multiple atlases further facilitates the use of this data by smaller labs and research groups with a more expansive range of atlases and tools. Additionally, CAROT may help with connectome-based meta analyses, for which there are few, by allowing results to be pooled across studies. Coordinate-based meta analyses are popular for task activation and brain morphometry studies (Laird et al., 2005; Yarkoni et al., 2011; Eickhoff et al., 2009) and are possible as most neuroimaging studies rely on a common template (*i.e*., the MNI template). This common template allows for spatial comparisons and pooling of results across different studies.

There are a few notable strengths and limitations of CAROT. First, CAROT appears to be robust to the choices of algorithmic parameters such as the number of fMRI frames, sample size used for training, the choice of cost matrix, and the equation used to solve the optimal transport problem. We showed that the method is not sensitive to parameter search in part due to the large amount of spatial and temporal autocorrelation in fMRI data (Shinn et al., 2021), which allow something as complex as a connectome to be compactly parameterized.

Future work includes generalizing CAROT to other functional time series data—such as electroencephalography (EEG), functional near infrared spectroscopy (fNIRS), or even wide-field CA2+ imaging data in mice (Lake et al., 2020)—where spatial and temporal autocorrelation patterns will be different. One limitation is that since CAROT is based on time series data, it is only appropriate for functional connectomes. Nevertheless, the ‘‘missing atlas” problem also exists for structural connectomes, for which no solution exists. Hence, the problem still exists for studies looking to uncover structure-function relationships at the connectome level. However, perhaps CAROT based on Euclidean distance rather than functional distance may be a reasonable approach to map between atlases used to create structural connectomes as well as map between different atlases used in morphometric analyses (such as the Desikan-Killiany and Destrieux atlases used in FreeSurfer). While we tested CAROT with an extensive range of atlases, we could not test CAROT in every functional atlas, as there are many. Nevertheless, given the range in atlas size (200-500 nodes) and atlas coverage (whole-brain and cortical only), we expect CAROT to work well for modern atlases not tested here and look to update CAROT when a new generation of brain atlases emerges.

In sum, CAROT allows a connectome generated based on one atlas to be directly transformed into a connectome based on another without needing raw data. These reconstructed connectomes are similar to and, in downstream analyses, behave like the original connectomes created from the raw data. Using CAROT on preprocessed open source data will increase its utility, accelerate the use of big data, and help make generalization and replication efforts easier.

## 6. Acknowledgments

This work was supported by NIMH R01 MH121095, NSF (IIS-1845032), ONR (N00014-19-1-2406), and Tata. Data were provided in part by the Human Connectome Project, WU-Minn Consortium (Principal Investigators: David Van Essen and Kamil Ugurbil; U54 MH091657) and funded by the 16 NIH Institutes and Centers that support the NIH Blueprint for Neuroscience Research; and by the McDonnell Center for Systems Neuroscience at Washington University. Data was also provided by the REST-meta-MDD Consortium Data Sharing, which was supported by the National Key R&D Program of China (2017YFC1309902), the National Natural Science Foundation of China (81671774, 81630031, 81471740 and 81371488), the 13th Five-year Informatization Plan (XXH13505) of Chinese Academy of Sciences, Beijing Municipal Science & Technology Commission (Z161100000216152, Z171100000117016, Z161100002616023 and Z171100000117012), Department of Science and Technology, Zhejiang Province (2015C03037) and the National Basic Research (973) Program (2015CB351702). The remainder of the data used in this study were provided by the Philadelphia Neurodevelopmental Cohort (Principal Investigators: Hakon Hakonarson and Raquel Gur; phs000607.v1.p1). Support for the collection of the data sets was provided by grant RC2MH089983 awarded to Raquel Gur and RC2MH089924 awarded to Hakon Hakonarson.

## Supplemental Materials

**Figure S1:**
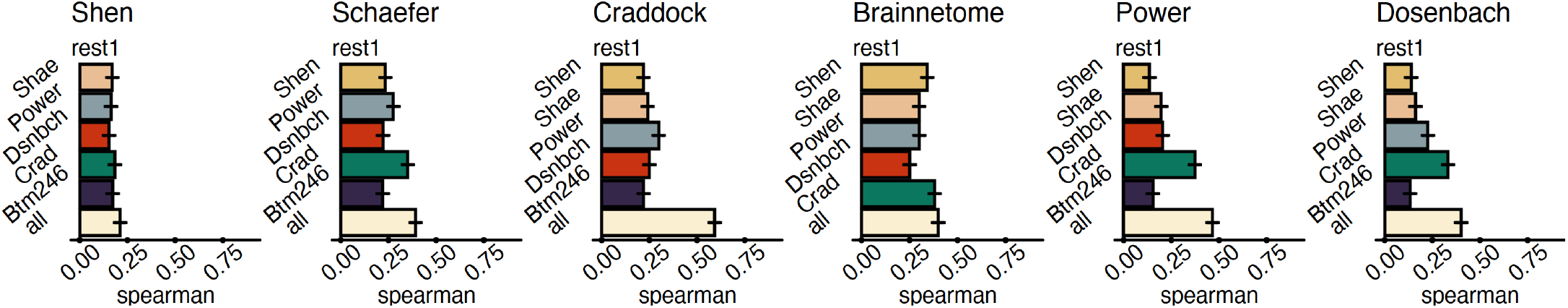
CAROT performance (rest) using euclidean distance between the center of gravity for each ROI as the cost measure. The results exhibit significantly lower performance compared to functional distance.

**Figure S2:**
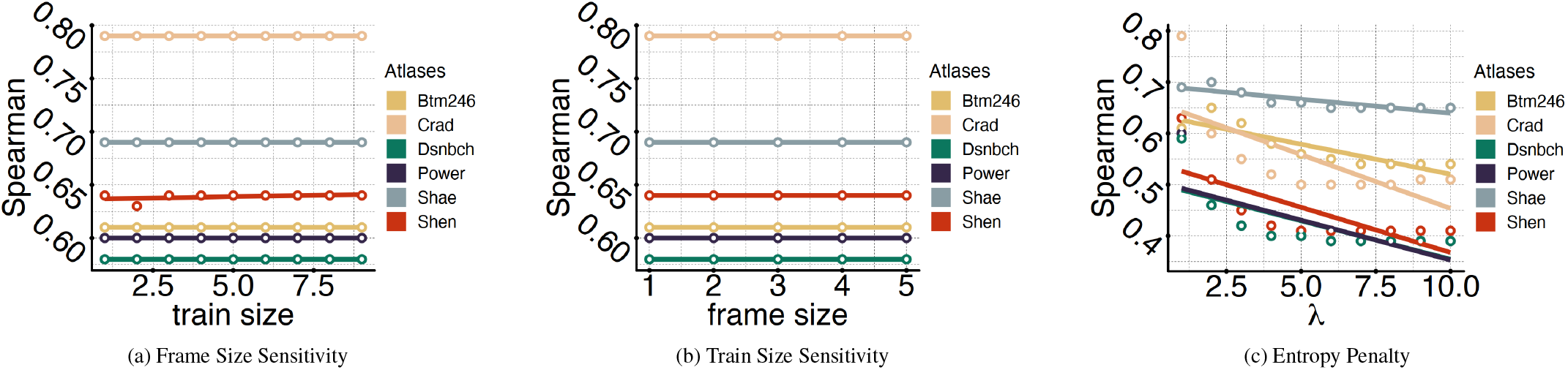
Parameter sensitivity of frame size, training data, and entropy regularization *ϵ* for different target atlases.

**Figure S3:**
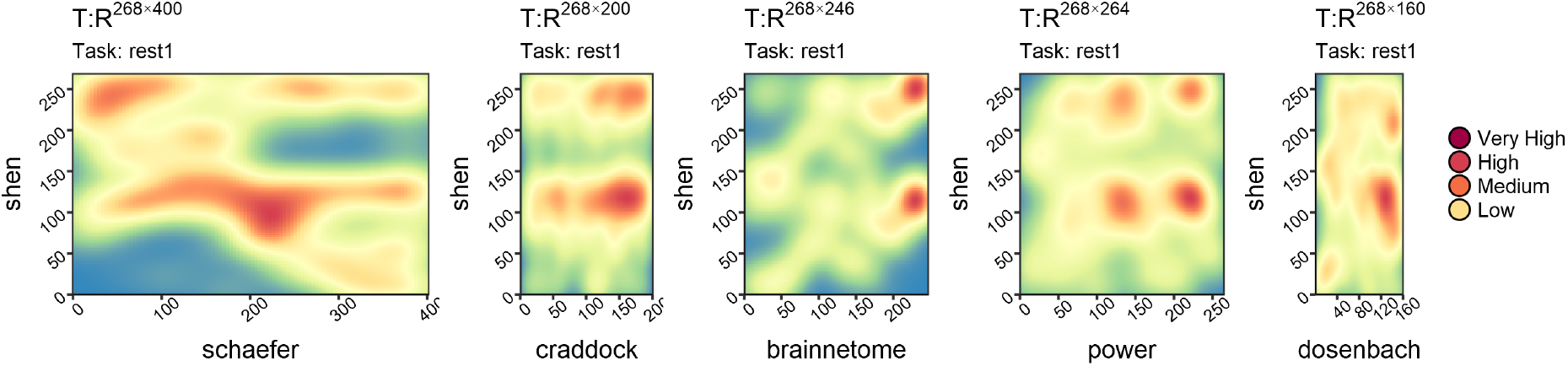
Optimal transport mappings derived from resting-resting and task data from the Shen atlas (source atlas) to each other atlas. Warmer color indicates regions that contribute the most towards mapping between atlases. Horizontal blue areas may indicate locations that are missing in the source atlas. For example, the Schaefer atlas does not include regions in the cerebellum, while the Shen does.

